# Rapid and Efficient Inactivation of SARS-CoV-2 from Surfaces using UVC Light Emitting Diode Device

**DOI:** 10.1101/2021.04.20.440654

**Authors:** Varun Dwivedi, Jun-Gyu Park, Stephen Grenon, Nicholas Medendorp, Cory Hallam, Jordi B. Torrelles, Luis Martinez-Sobrido, Viraj Kulkarni

## Abstract

Efforts are underway to develop countermeasures to prevent the environmental spread of COVID-19 pandemic caused by SARS-CoV-2. Physical decontamination methods like Ultraviolet radiation has shown to be promising. Here, we describe a novel device emitting ultraviolet C radiation (UVC), called NuvaWave, to rapidly and efficiently inactivate SARS-CoV-2. SARS-CoV-2 was dried on a chambered glass slides and introduced in a NuvaWave robotic testing unit. The robot simulated waving NuvaWave over the virus at a pre-determined UVC radiation dose of 1, 2, 4 and 8 seconds. Post-UVC exposure, virus was recovered and titered by plaque assay in Vero E6 cells. We observed that relative control (no UVC exposure), exposure of the virus to UVC for one or two seconds resulted in a >2.9 and 3.8 log_10_ reduction in viral titers, respectively. Exposure of the virus to UVC for four or eight seconds resulted in a reduction of greater than 4.7-log_10_ reduction in viral titers. The NuvaWave device inactivates SARS-CoV-2 on surfaces to below the limit of detection within one to four seconds of UVC irradiation. This device can be deployed to rapidly disinfect surfaces from SARS-CoV-2, and to assist in mitigating its spread in a variety of settings.

## Introduction

The coronavirus disease 2019 (COVID-19) pandemic caused by severe acute respiratory syndrome-coronavirus-2 (SARS-CoV-2) has caused a major public health and economic crisis.^1^ It is reported that SARS-CoV-2 originated in Wuhan city, China’s Hebei province, in December 2019.^2^ This virus spread rapidly all over the world, prompting the World Health Organization (WHO) to declare COVID-19 a pandemic in March 2020.^3^ As of this writing, there are over 93 million cases of SARS-CoV-2 infection and more than 2 million deaths due to COVID-19 worldwide.^4^ Urgent mitigation efforts are needed to control this pandemic. Several prophylactics and therapeutics are undergoing human clinical trials.^5^ Currently, the United States Food and Drug Administration (US-FDA) has approved one therapeutic antiviral drug, Remdesivir, and a monoclonal antibody, MAb (Bamlanivimab) the treatment of SARS-CoV-2 infections under the Emergency Use Authorization (EUA) program.^6^ Recently, two mRNA-based vaccines developed by Pfizer-BioNTech and Moderna have been approved under the EUA by the US-FDA.^7, 8^

Person-to-person transmission of the SARS-CoV-2 is primarily via aerosols expelled by an infectious person and inhales via a susceptible person.^9, 10^ SARS-CoV-2 transmission through contaminated surfaces has also been reported.^10^ Super spreader events have occurred at places where there are large public gatherings, including malls, restaurants, airports, schools, train and bus stations, and hospitals, among others. Respiratory viruses including SARS-CoV-2 can survive for hours to days on surfaces.^11-13^ Several studies have demonstrated SARS-CoV-2 RNA in healthcare and community settings such as house-holds, day care centers, schools, and airports.^11, 14-17^ Thus, effective environmental disinfection of public locations is needed to mitigate the transmission of SARS-CoV-2 and control the COVID-19 pandemic.

Germicidal ultraviolet C (UVC) irradiation has proven to be effective to inactivate pathogens from air and surfaces in public areas. ^18-23^ Here we describe a next generation NuvaWave device that generate UVC germicidal irradiation. NuvaWave is a powerful, lightweight, intelligent, UVC handheld device operating at 270 nm, near the peak of germicidal efficacy. It is intended to disinfect surfaces with non-ionizing UVC irradiation. In this study, NuvaWave UVC light was tested for its virucidal activity against SARS-CoV-2. Our results indicate that the UVC light generated from NuvaWave is highly efficient in inactivating SARS-CoV-2 in under 4 seconds. Our results support the feasibility of using NuvaWave to rapidly disinfect surfaces from SARS-CoV-2 to prevent viral transmission from surfaces to humans in an effort to interrupt SARS-CoV-2 transmission and mitigate the COVID-19 pandemic.

## MATERIAL AND METHODS

### Cell line and virus

Vero E6 cells (CRL-1586) were procured from the American Type Collection (ATCC, Manassas, VA). Cells were maintained in Dulbecco′s Modified Eagle′s Medium (DMEM, Corning, Glendale, AZ) containing 10% fetal bovine serum (FBS), L-glutamine, 1% penicillin, streptomycin and L-glutamine (PSG, Corning, Glendale, AZ). Vero E6 cells were used to generate SARS-CoV-2 stocks and for virus titration assays. SARS-CoV-2, USA-WA1/2020 strain, passage 4 (P4) (Gen Bank MN985325.1) was obtained from the Biodefense and Emerging Infections (BEI, Manassas, VA) Research Resources Repository (catalog number NR-52281). Virus stocks were generated by passaging SARS-CoV-2 in Vero E6 cells at a multiplicity of infection (MOI) of 0.001. This master virus stock (passage 5, P5) was used to generate a sixth cell culture passage stock (P6) by infecting Vero E6 cells at a MOI of 0.02. P6 viral stock was used to generate a P7 working stock. The resulting viral P7 stock had a titer of 3.5 x 10^6^ plaque forming units (PFU)/ml and was used to test the efficacy of NuvaWave UVC device. All work was carried out in the state-of-art biosafety level 3 (BSL3) laboratories at Texas Biomedical Research Institute with the approval of the Institutional Biohazard Committee.

### NuvaWave UVC device

NuvaWave is a powerful, lightweight, computer controlled, narrowband UVC handheld device operating at 270 nm, near the peak of germicidal efficacy (**Fig. 1a**). It is intended to disinfect surfaces with non-ionizing UVC radiation by waving the device 1 to 3.5 inches over the surface. The patent-pending technology ensures adequate germicidal dosage is delivered over the entire 4 inches x 4 inches exposure area and at distances between 1 and 3.5 inches. The UVC light source is instant on/off and is controlled with a simple trigger mechanism. The system utilizes an external battery pack rated for up to 3 hours of use and incorporates a computer monitoring system which ensures consistent performance over time. To test the efficacy of this device the same light engine as the NuvaWave Handheld and packaged in a computer controlled robotic test fixture (**Fig. 1b**). The robotic test fixture was engineered to hold a chambered glass slide on to which virus sample was placed.

**Figure 1.**
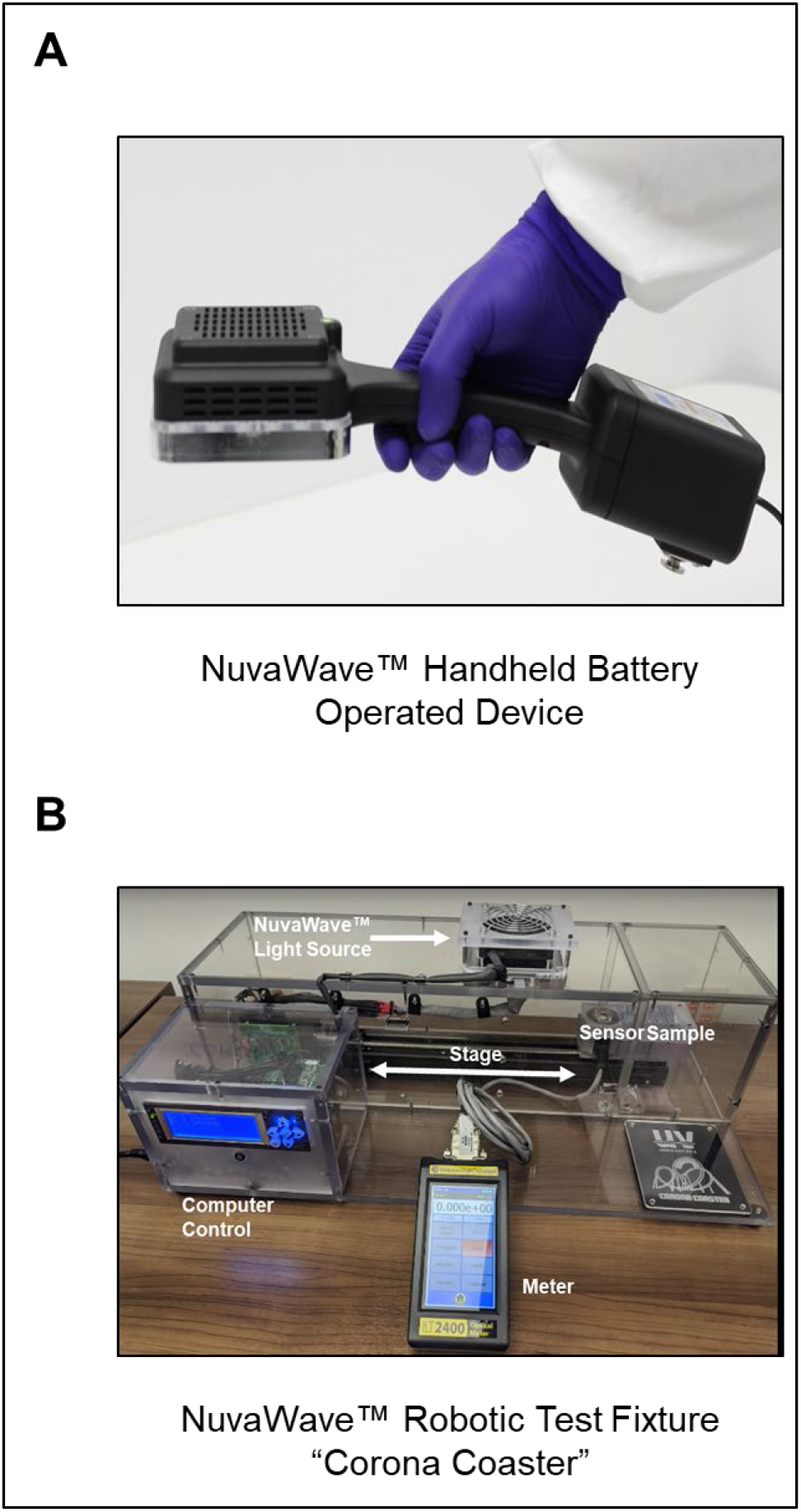
NuvaWave Handheld Battery-Operated Device and robotic test fixture. (A) NuvaWave is a powerful, lightweight, computer controlled, UVC handheld device operating at 270 nm, near the peak of germicidal efficacy. It is intended to disinfect surfaces with non-ionizing UVC radiation by waving the device 1 to 3.5 inches over the surface. (B) A specialized test fixture to mimic waving the handheld over a surface at a constant speed for a single pass. This unit was used to test the effect of UVC on SARS-CoV-2.

### Exposure of SARS-CoV-2 to UVC

Fifty microliters containing 3.5 x 10^6^ PFU/ml SARS-CoV-2 viral stock was placed on one well of a 4-well chambered glass slide (Nunc, Sigma) (**Fig 2**). Five slides were used for 5 different exposure conditions: no exposure, 1, 2, 4 and 8 seconds UVC transverse exposure times. The virus was allowed to dry in a biosafety cabinet in a BSL3 laboratory for 1 hour at room temperature (RT). After drying, the slide was placed into the NuvaWave Robotic Test Fixture and exposed to UVC light radiation for the predetermined exposure conditions. Immediately after exposure the virus was reconstituted in 50 ml of DMEM and 10-fold serial dilutions were performed to measure viral titers.

**Figure 2:**
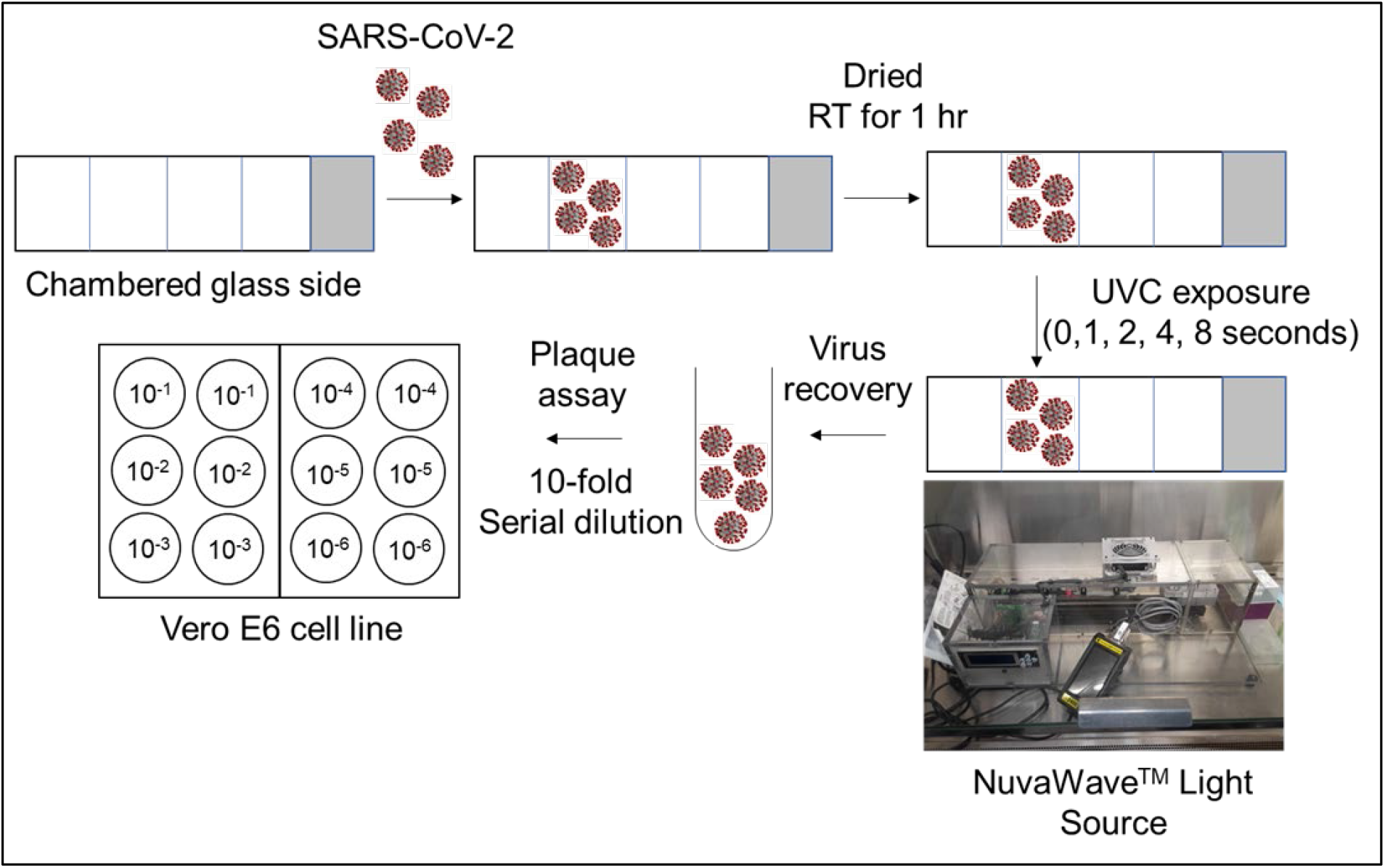
Schematic representation of the workflow. SARS-CoV-2 was placed on the chambered glass slide, dried for 1 hour at room temperature, and then exposed to UVC light radiation using the NuvaWave device for 1, 2, 4 and 8 seconds. No UVC light exposure was used as control. After UVC light exposure, virus was recovered, serial diluted and used to assess viable virus in Vero E6 cells by the plaque assay.

### Viral titrations

Viral titers were determined by plaque assay. Briefly, confluent monolayers of Vero E6 cells (12-well format, 0.5 x 10^5^ cells/well, duplicates) were infected with 10-fold serial dilutions of the reconstituted virus for 1 hour at 37°C in a humidified 5% CO_2_ incubator. Following virus adsorption, cells were washed and overlaid with post-infection media containing 2% agar (Oxoid, Hampshire, UK). Plates were incubated for 48 hours at 37°C in a humidified 5% CO_2_ incubator. After 48 hours, plates were submerged in 10% neutral buffered formalin (Sigma-Aldrich, St. Louis, MO) for 24 hours to fix cells and inactivate the virus. Presence of virus was evaluated by immunostaining. Briefly, cells were washed three times with PBS and permeabilized with 0.5% Triton X-100 (Sigma-Aldrich, St. Louis, MO) for 10 minutes at RT. Cells were then blocked with 2.5% bovine serum albumin (BSA) (Sigma-Aldrich, St. Louis, MO) in PBS for 1 hour at 37°C, followed by incubation with 1 µg/ml of a SARS-CoV nucleocapsid protein (NP) monoclonal antibody (mAb) 1C7C7 (Millipore-Sigma, St. Louis, MO) shown to cross-react with SARS-CoV-2 NP, diluted in 1% BSA for 1 hour at 37°C.^24^ After incubation with the primary mAb, cells were washed three times with PBS, and developed with the Vectastain ABC kit and DAB Peroxidase Substrate kit (Vector Laboratory, Inc., Burlingame CA) according to the manufacturers’ instructions.^24^ Viral titers were calculated as PFU/mL.

### Statistics

Data obtained was plotted using the GraphPad Prism v.8.0 software (GraphPad Software, San Diego, CA) and significance determined by one-way ANOVA Tukey post-test comparing experimental groups over time.

## RESULTS

### Rapid and efficient inactivation of SARS-CoV-2

The NuvaWave Light Source is comprised of 32 UVC light-emitting diodes (LEDs) centered at 270 nm. Individual reflectors for each LED have been designed to provide uniform power distribution across a radiation area of 4 inches x 4 inches and a depth between 1 inch and 3.5 inches. The NuvaWave handheld device is shown in **Fig. 1A**. Testing was performed using the same light engine as the NuvaWave Handheld and packaged in a computer controlled robotic test fixture (**Fig. 1B**). The light engine was mounted 2 inches from the sample while the sample and a sensor were moved on a stage at a constant speed under the light source. A National Institute of Standards and Technology (NIST) calibrated sensor (International Light Technologies model SED270) and meter (International Light Technologies model ILT2400) were used to measure the precise exposure dosage in Joules/cm^2^. We tested the ability of the UVC light generated by the NuvaWave UVC device on SARS-CoV-2 inactivation following the workflow schematic depicted in **Fig. 2**.

In this study, we mimic the NuvaWave operator waving the device in a single pass over a surface. To facilitate the study, a robotic testing system was created (**Fig. 1B**) in which we applied the illumination engine on top of a test chamber and automated the movement of SARS-CoV-2 at a controlled rate across the UVC light source aperture while maintaining a 2-inch distance. The NuvaWave Robotic Tester includes an integrated sensor, that moved with the SARS-CoV-2, to measure the dosage (J/cm^2^) that the SARS-CoV-2 was exposed during the irradiation test. Viral samples traveled at constant speed across the entire 4-inch UVC illumination aperture. The transverse time is defined as the length of time it takes for the sample to move through the 4 inches UVC illumination aperture. The effectiveness was evaluated at four transverse times of 1, 2, 4, and 8 seconds. This tester was built with a polycarbonate surrounding, so the operator was not exposed to any stray UV light (**Fig. 1B**). This device is identical to the commercially available product both in the light source design and the electronics driving the light source. It differs in packaging only. New packaging was needed to incorporate the stage and allow the Robotic Test Fixture to enable and disable the light source without human intervention thus minimizing any human error in the UVC exposure methodology. UVC dosage (Joules/cm^2^) were recorded after each exposure (**Fig. 3**). The dosage of UVC recorded from 5 individual repeated experiments were consistent, showing the reliability of the instrument tested in the accurate deployment of the UVC dosage. The exposures for 1, 2, 4, 8 seconds were 0.012 J/cm^2^, 0.025 J/cm^2^, 0.050 J/cm^2^, 0.100 J/cm^2^ respectively (**Fig. 3)**.

**Figure 3:**
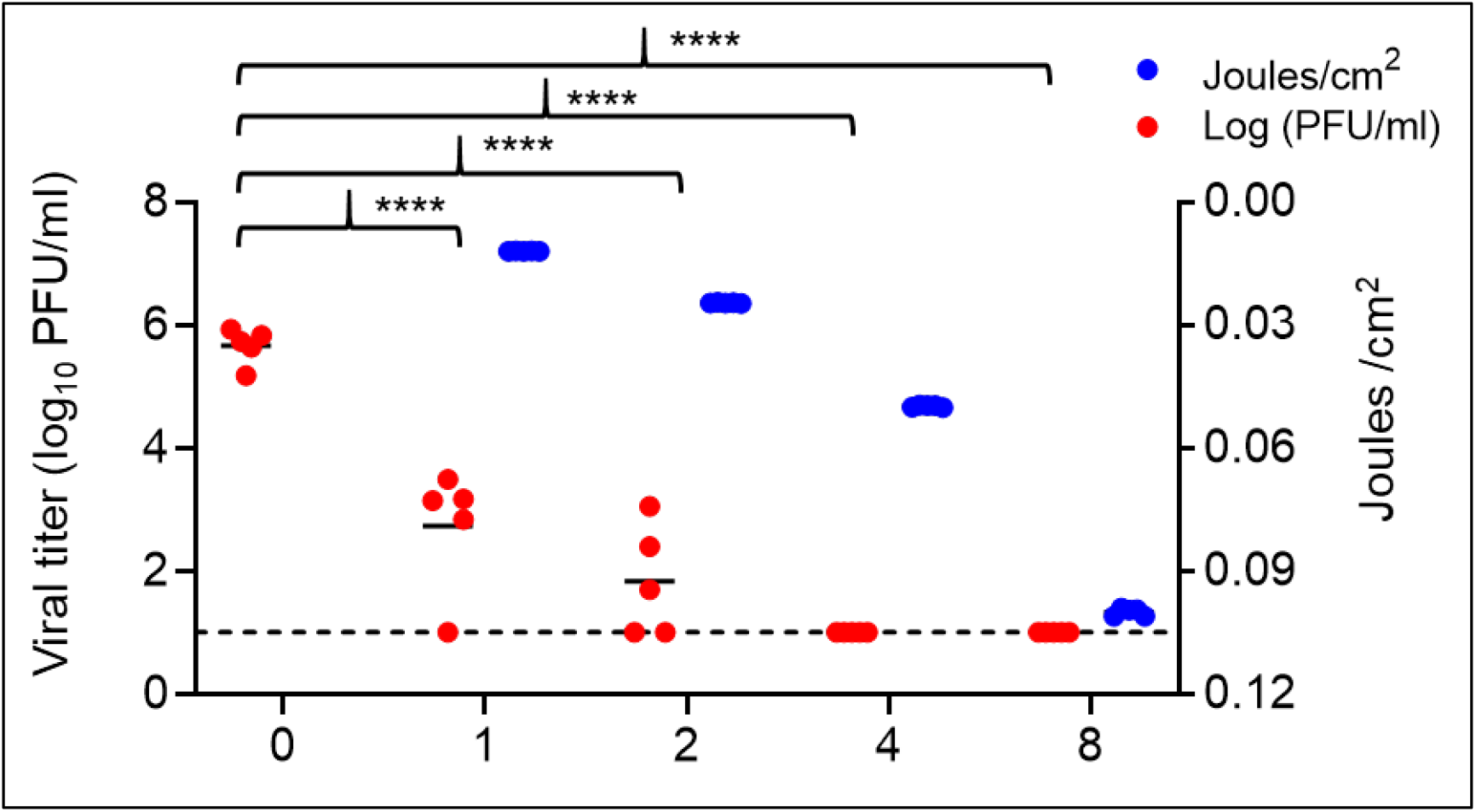
Efficacy of NuvaWave UVC device against SARS-CoV-2. NuvaWave UVC device was tested against SARS-CoV-2 in five independent experiments (n=5, Exp 1-5). Dosage of UVC radiation after 1, 2, 4 and 8 second exposure time (blue circles, right Y-axis). SARS-CoV-2 titers as determined by plaque assay in Vero E6 cells (red circles, left Y-axis). Viral titers (PFU/ml) under no exposure (control) and 1, 2, 4 and 8 seconds UVC radiation exposure are shown. Dotted line indicates the limit of detection (LOD, 10 PFU). Viral titers from no UVC exposure were compared to viral titers obtained after UVC exposure at indicated time points (seconds); p values were calculated by One-way ANOVA post-Tukey test by using GraphPad Prism vr.8 software.

For our unexposed controls, we report a viral recovery of 5.7 log_10_ plaque forming units (mean value of n=5 in duplicate) per ml (PFU/ml) (**Fig. 3** and **Table 1)**. UVC light exposed virus for one or two seconds resulted in >2.9 log_10_ and >3.8 log_10_ reductions in viral titers, respectively, when compared to the unexposed control (**Fig. 3** and **Table 1)**. Exposure of the virus to UVC for four or eight seconds resulted in both cases in a reduction of over the detection limit of the assay (>4.7 log_10_ reduction) in viral titers relative to our unexposed control (**Fig. 3** and **Table 1**). Viral titers were statistically lower in all UVC light exposed virus for all the time points tested (1 to 8 seconds) when compared to unexposed controls (*p*<0.0001) (**Fig. 3**). Importantly, the average UVC dosage of 0.05 J/cm^2^ obtained at 4 second exposure time, resulted in >99.998% reduction in SARS-CoV-2 (>4.7 log_10_ reduction). We observed progressive reduction in viral titers as the dosage of UVC increased. These results indicate that UVC light radiation generated by NuvaWave device inactivates SARS-CoV-2 from glass surfaces under our defined experimental conditions (**Table 1**) suggesting the feasibility of using NuvaWave UVC device for the decontamination of surfaces from SARS-CoV-2.

**Table 1:** Virucidal activity of UVC radiation generated by the NuvaWave device.

| Time<br>(seconds) | Viral Titer<br>PFU/ml (log<sub>10</sub>) |  |  |  |  |  | Reduction |  |
| --- | --- | --- | --- | --- | --- | --- | --- | --- |
|  | Exp 1 | Exp 2 | Exp 3 | Exp 4 | Exp 5 | Avg | % | log<sub>10</sub> |
| 0 | 5.64 | 5.18 | 5.93 | 5.83 | 5.73 | 5.7 | 0 | 0 |
| 1 | <1 | 3.17 | 3.49 | 2.84 | 3.14 | 2.7* | >99.884% | >2.9 |
| 2 | <1 | 3.05 | <1 | 2.4 | 1.7 | 1.8* | >99.985% | >3.8 |
| 4 | <1 | <1 | <1 | <1 | <1 | <1* | >99.998% | >4.7 |
| 8 | <1 | <1 | <1 | <1 | <1 | <1* | >99.998% | >4.7 |

Log<sub>10</sub> and percentage reduction in viral titers after exposure to UVC radiation at indicated time (seconds) is reported. Data from 5 independent experiments are shown.
PFU, plaque forming units;
Exp, experiment
Avg, average
Note: <1-log<sub>10</sub> = below the limit of detection; \*<1-log<sub>10</sub> was considered as 1-log<sub>10</sub> for calculation purposes

The UVC light irradiation at a distance of 2 inches can consistently inactivate SARS-CoV-2 placed on a hard glass surface within a few seconds. To the best of our knowledge, this is the first report on UVC light that can inactivate SARS-CoV-2 within seconds of light exposure at the given experimental conditions.

## DISCUSSION

SARS-CoV-2 is a novel coronavirus identified in 2019 and the causative agent of the COVID-19 pandemic. ^3, 9^ The primary mode of spread of SARS-CoV-2 is via respiratory aerosols.^1, 9, 25^ Effective countermeasures are necessary to stop the spread of SARS-CoV-2 to control the ongoing COVID-19 pandemic. In addition to the development of effective vaccines and antivirals, there is also an urgent need for effective disinfection methods to inactivate the virus on environmental surfaces. Aerosol transmission and exposure to contaminated surfaces at public places play a major role in the spread of SARS-CoV-2.^13^ UVC irradiation has been shown to be effective against a number of human pathogens.^18-20, 26-28^ SARS-CoV-2 has been reported to be inactivated by UVC irradiation.^18, 28, 29^ However, some devices that generate UVC radiation have required prolonged periods of exposure to inactivate SARS-CoV-2. Indeed, the use of far-UVC light (222 nm) to inactivate airborne human coronaviruses with 99.9% efficacy in ∼25 minutes of light exposure;^18^ and the use of a pulsed-xenon, multispectral UVC light to inactivate SARS-CoV-2 from glass surfaces and N95 respirators have been reported. In the latter study, authors observed 3.54 log_10_, >4.54 log_10_ and >4.12 log_10_ viral reduction when exposed to UVC light for 1, 2 and 5 minutes respectively.^30^ Here, we describe a device that emits UVC that inactivates SARS-CoV-2 in less than 4 seconds.

We tested the NuvaWave UVC device to determine the efficiency of its generated UVC light in inactivating SARS-CoV-2 on a glass surface. The main characteristics of a glass surface are the transparency, heat resistance, pressure and breakage resistance and chemical resistance. At both 4 and 8 second UVC exposures, we observed >4.7 log_10_ reduction compared to our control (UVC unexposed virus). Whereas, at 1 and 2 second UVC exposures, we observed >2.9 log_10_ and >3.8 log_10_ virus reduction, respectively. Thus, the UVC radiation from NuvaWave device is effective in inactivating SARS-CoV-2. Overall, our findings show that NuvaWave UVC device is able to inactivate SARS-CoV-2 from surfaces within seconds. We have not tested UVC radiation from this device on inactivation of SARS-CoV-2 on other materials. However, based on our findings, we anticipate that this device will inactivate SARS-CoV-2 on other hard surfaces and fomites, and thus, NuvaWave device could be deployed to mitigate the environmental spread of SARS-CoV-2. Likewise, we are currently assessing the ability of the NuvaWave UVC device to inactivate other human pathogens.

## Acknowledgements

We would like to thank the Texas Biomedical Research Institute BSL-3 Operations Program, the Environmental Health Service, and the Texas Biomedical Research Institute Biosafety Committee for their support in carrying out this study.

## Author contributions

VK, VD, LMS, JBT, NM designed the experiments; VK and VD performed all experiments, gathered and analyzed data; SG, NM designed the NuvaWave device; VK, VD, LMS, JBT, SG, NM and CH reviewed data; VK wrote the manuscript with inputs and revisions from all other co-authors. All authors approved the manuscript.

